# Genetic disruption of dopamine β-hydroxylase dysregulates innate responses to predator odor in mice

**DOI:** 10.1101/2023.06.21.545975

**Authors:** Joyce Liu, Daniel J. Lustberg, Abigail Galvez, L. Cameron Liles, Katharine E. McCann, David Weinshenker

## Abstract

In rodents, exposure to predator odors such as cat urine acts as a severe stressor that engages innate defensive behaviors critical for survival in the wild. The neurotransmitters norepinephrine (NE) and dopamine (DA) modulate anxiety and predator odor responses, and we have shown previously that dopamine β-hydroxylase knockout (*Dbh -/-)*, which reduces NE and increases DA in mouse noradrenergic neurons, disrupts innate behaviors in response to mild stressors such as novelty. We examined the consequences of *Dbh* knockout (*Dbh -/-*) on responses to predator odor (bobcat urine) and compared them to Dbh-competent littermate controls. Over the first 10 min of predator odor exposure, controls exhibited robust defensive burying behavior, whereas *Dbh -/-* mice showed high levels of grooming. Defensive burying was potently suppressed in controls by drugs that reduce NE transmission, while excessive grooming in *Dbh -/-* mice was blocked by DA receptor antagonism. In response to a cotton square scented with a novel “neutral” odor (lavender), most control mice shredded the material, built a nest, and fell asleep within 90 min. *Dbh -/-* mice failed to shred the lavender-scented nestlet, but still fell asleep. In contrast, controls sustained high levels of arousal throughout the predator odor test and did not build nests, while *Dbh -/-* mice were asleep by the 90-min time point, often in shredded bobcat urine-soaked nesting material. Compared with controls exposed to predator odor, *Dbh -/-* mice demonstrated decreased c-fos induction in the anterior cingulate cortex, lateral septum, periaqueductal gray, and bed nucleus of the stria terminalis, but increased c-fos in the locus coeruleus and medial amygdala. These data indicate that relative ratios of central NE and DA signaling coordinate the type and valence of responses to predator odor.

## 1. Introduction

### 1.1. Innate fear responses

Responding to external stimuli with appropriate behaviors (e.g., approach or avoidance) is crucial for survival of most animals. In rodents, detection of cat odorants prompts adaptive defensive and escape behaviors that reduce risk of predation. It is important to understand the neurobiological underpinnings of species-typical and maladaptive responses to threats because dysregulated behavioral responses to stressful stimuli represent many hallmark features of neuropsychiatric disorders, including posttraumatic stress disorder (PTSD), anxiety, and substance use disorder (Dias et al 2013).

### 1.2 Norepinephrine modulates anxiety-like traits

The catecholamine neuromodulators norepinephrine (NE) and dopamine (DA) govern emotional and motivated behavior. NE is critical for arousal, attention, orienting to salient stimuli, and organization of stress responses (Poe et al 2020, Sara 2009), while DA mediates reward and approach (Krach et al 2010). The locus coeruleus (LC) is the main source of NE throughout the brain and projects to regions implicated in responses to potential threats (Lustberg et al 2020a, Lustberg et al 2020b, Poe et al 2020, Schwarz & Luo 2015). Although NE is the dominant neurotransmitter produced by the LC, it can also release DA under certain conditions (Beas et al 2018, Kempadoo et al 2016, Petter et al 2023, Takeuchi et al 2016).

The LC is activated by novelty and predator odor, and NE transmission facilitates anxiety-like behavior, particularly in paradigms that involve stress and novelty (Curtis et al 2012, Day et al 2004, Hayley et al 2001, Lustberg et al 2020b). NE has also been previously implicated in predator odor responses as well. Exposure to trimethylthiazoline (TMT), a component of fox odor, promotes freezing behavior in rats, which is attenuated by pharmacological blockade of NE transmission in the bed nucleus of the stria terminalis (BNST) (Fendt et al 2005), and systemic or intra-dorsal premammilary nucleus β-adrenergic receptor antagonism abates defensive responses to cat odor (De Monte et al 2008). Likewise, CRISPR-induced disruption of NE synthesis in the mouse LC blocks the ability of rat odor to trigger arousal and wakefulness (Yamaguchi et al 2018).

### 1.3 Norepinephrine and dopamine control distinct stress-induced behaviors

We have previously used dopamine β-hydroxylase knockout (*Dbh -/-*) mice that lack NE and have excessive DA in LC and other noradrenergic cell groups to investigate the relative contribution of these catecholamines to stress responses. The anxiogenic effects of novelty stress are abolished in *Dbh -/-* mice, and they also fail to exhibit species-typical repetitive burying and nestlet shredding behaviors in novel environments (Lustberg et al 2020a, Lustberg et al 2020b). Pharmacological restoration of NE levels (without normalizing DA levels) rescues these phenotypes, while pharmacological blockade of NE transmission recapitulates them in normal mice. These findings indicate that the lack of NE is primarily responsible for the aberrant responses.

However, we recently reported that both NE and DA contribute to novel odorant stress (Lustberg et al 2022). In response to non-threatening but novel olfactory stimuli (e.g. plant-derived essential oils presented in a novel environment), normal mice escalate digging, burrowing, and burying. In contrast, *Dbh -/-* mice do not engage in these behaviors, and instead groom excessively in novel environments regardless of the presence of various “neutral” odors. Importantly, acute suppression of NE transmission blocks novel odor-induced digging in control mice, while DA receptor antagonists attenuate grooming in *Dbh-/-* mice, indicating that the NE:DA ratio in noradrenergic neurons modulates the nature of these behavioral responses.

### 1.4 Study objectives and summary

While our previous work has provided insight concerning the roles of NE and DA in behavioral and brain responses to mild stressors such as novel environments and novel (but non-threatening) odors, the contribution of these catecholamines to severe innate threats has not been fully identified. Because loss of DBH uniquely decreases NE while simultaneous increasing DA, we speculated that *Dbh -/-* mice would lack species-typical defensive responses to predator odor and potentially display paradoxical appetitive/approach behaviors towards it. To test this, we exposed *Dbh -/-* and control mice to neutral odors or bobcat urine applied to cotton nestlets in a novel environment and measured defensive burying, grooming, nestlet shredding, and arousal. We then used a pharmacological approach to dissect the relative contributions of individual NE and DA receptor subtypes. We also assessed the effects of DBH knockout on molecular signatures of neuronal activation in brain regions that receive innervation from the LC and are implicated in behavioral responses to predator odor.

## 2. Materials and methods

### 2.1 Animals

A total of 40 *Dbh -/-* mice, maintained on a mixed 129/SvEv and C57BL/6 J background as previously described (Thomas et al 1998, Thomas et al 1995), were used in this study. *Dbh -/-* males were bred to *Dbh +/-* females, and pregnant *Dbh* +/- dams were administered drinking water containing the β-adrenergic receptor (AR) agonist isoproterenol and the α1AR agonist phenylephrine (20 μg/ml each; Sigma-Aldrich) with vitamin C (2 mg/ml) from E9.5– E14.5, and the synthetic NE precursor L-3,4-dihydroxyphenylserine (DOPS; 2 mg/ml; Lundbeck, Deerfield, IL) + vitamin C (2 mg/ml) from E14.5-parturition to prevent embryonic lethality resulting from complete *Dbh* deficiency. *Dbh* -/- mice are easily distinguished from their NE-competent littermates by their visible delayed growth and bilateral ptosis phenotypes. A total of 28 *Dbh* +/− littermates were used as controls because their behavior and catecholamine levels are indistinguishable from wild-type (*Dbh +/+*) mice (Bourdelat-Parks et al 2005, Szot et al 1999, Thomas et al 1998). Adult mice 4-10 months of age were used for all experiments.

Because no sex differences in stress-induced digging or grooming have been reported in past literature (Dixit et al 2020, Londei et al 1998, Smolinsky AN 2009) or were observed in published experiments from our lab (Lustberg et al 2022) and pilot experiments from this study, male and female mice from the same *Dbh* genotype were evenly distributed across drug treatment groups, and data were pooled between sexes. All animal procedures and protocols were approved by the Emory University Animal Care and Use Committee, in accordance with the National Institutes of Health guidelines for the care and use of laboratory animals. Mice were maintained on a 12 h/12 h light/dark cycle (7:00/19:00 h) with access to food and water ad libitum, except during behavioral testing. Behavioral testing was conducted under standard lighting conditions during the light cycle (14:00-17:00) in the same room where the mice were housed to minimize the stress of cage transport on test days.

### 2.2 Behavioral analysis

Mice were group-housed in static cages in same-sex groups of 2–5. In the first set of experiments, individual mice were removed from their home cages and placed into new standard mouse cages (13″ x 7″ x 6″) containing only clean bedding substrate and a cotton nestlet square (2” × 2”, ∼3 g) pre-soaked with either 1 ml of deionized water (odorless control) or predator odor (bobcat urine; Maine Outdoor Solutions, LLC, Hermon, Maine). Each mouse received both treatments in a counterbalanced order spaced 2 d apart. A clear plexiglass cover was pressed flush over the cages to prevent vaporization or dispersal of the odorants over the course of testing, while still allowing filming and behavioral scoring. The mice in the odorant-treated cages were filmed with a front-facing, mounted digital video camera for 10 min for assessment of defensive (digging/burying) and non-defensive (grooming) behaviors (De Boer & Koolhaas 2003, Kalueff & Tuohimaa 2005, Kemble & Bolwahnn 1997, Londei et al 1998, Lustberg et al 2020a, Pond et al 2021, Smolinsky AN 2009). Time spent in defensive burying and grooming was manually scored with digital stopwatches by a trained observer blind to genotype and treatment. Defensive burying involved vigorous displacement of the bedding toward the odorant stimulus, often resulting in the burying of the odorant-soaked nestlet. Defensive burying sometimes presented as intense burrowing away from the stimulus, which resembled “swimming” or tunneling through the bedding substrate (Bondi et al 2007, De Boer & Koolhaas 2003, Fucich & Morilak 2018, Lustberg et al 2020a, Treit et al 1981). Burying is considered aversion-related, because it resembles responses exhibited after exposure to the shock probe test or predator odor (Garbe CM 1993, Hwa et al 2019, McGregor et al 2002, Sluyter et al 1996). Grooming was defined as repetitive paw movements oriented around the whiskers and face, as well as licking or scratching of the body and tail (De Boer & Koolhaas 2003, Kalueff & Tuohimaa 2005, Smolinsky AN 2009). Some reports suggest that grooming may be a self-soothing behavior that increases under stressful conditions, while others propose that it is a “rest behavior’ that indicates low levels of stress (Guild & Dunn 1982, Kemble & Bolwahnn 1997, Kemble ED 1995). For experiments with D1 receptor antagonist (see below), locomotor activity was measured as distanced traveled (m) using ANY-maze software (version 6.0, Stoelting Co, Wood Dale, IL, USA).

The second set of experiments was identical to the first set of experiments, except that mice were pretreated with vehicle, adrenergic, or dopaminergic drugs (see below) prior to predator odor exposure. All drugs were administered 30 min prior to testing except nepicastat, which was administered 2 h prior to testing. Each mouse received both a vehicle and drug pretreatment in a counterbalanced order spaced 2 d apart. Each mouse received no more than two different drugs, and trials with the same mice were spaced at least two weeks apart.

The third set of experiments were identical to the first set of experiments except that (1) the cotton nestlet square pre-soaked with either 1 ml of a novel “neutral” odor (lavender oil, 1:100 dilution in deionized water; Anjou, Fremont, CA) or bobcat urine, and (2) mice were left undisturbed for 90 min, after which a trained experimenter returned to record whether the mouse was asleep and whether the nestlet was fully (100%) shredded. Sleep was assessed behaviorally using EEG-validated scoring criteria, as we have described (Porter-Stransky et al 2019).

### 2.3 Drugs

The following drugs were intraperitoneally administered to dissect the relative contributions of NE and DA receptor signaling to behavior: nepicastat (100 mg/kg, DBH inhibitor) (Synosia Therapeutics, Basel, Switzerland); prazosin (0.5 mg/kg, α1AR antagonist) (Sigma-Aldrich, St. Louis, MO); propranolol (5 mg/kg, βAR antagonist) (Sigma-Aldrich); atipamezole (1 mg/kg, α2AR antagonist) (Sigma-Aldrich), flupenthixol (0.25 mg/kg, nonspecific DA receptor antagonist) (Sigma-Aldrich); SCH-23390 (0.03) mg/kg, D1 receptor antagonist) (Sigma-Aldrich); L-741,626 (10 mg/kg, D2 receptor antagonist) (Sigma-Aldrich). Doses selected were based on previous studies and pilot experiments to control for confounding effects such as motor impairment (Lustberg et al 2020a, Lustberg et al 2022, Mitchell et al 2008, Pina & Cunningham 2014, Yan et al 2016). Because the 0.03 mg/kg dose of SCH-23390 suppressed general motor activity, we also tested 0.01 and 0.006 mg/kg doses. All drugs were dissolved in bacteriostatic saline except prazosin and L-741,626. Prazosin was first dissolved in 1.5% DMSO and 1.5% Cremophor EL before being added to saline, and L-741,626 was first dissolved in 10% ethanol and 1.5% Cremophor EL before being added to saline. The corresponding vehicle that each drug was dissolved in was used as a control.

### 2.4 c-fos immunohistochemistry (IHC)

*Dbh +/-* and *Dbh -/-* mice were exposed to predator odor for 90 min as described above, then euthanized with an overdose of sodium pentobarbital (Fatal Plus, 150 mg/kg, i.p.; Med-Vet International, Mettawa, IL) for transcardial perfusion with cold 4% paraformaldehyde in 0.01 M PBS. After extraction, brains were post-fixed for 24 h in 4% paraformaldehyde at 4°C, and then transferred to cryoprotectant 30% sucrose/PBS solution for 72 h at 4°C. Brains were embedded in OCT medium (Tissue-Tek; Sakura, Torrance, CA) and serially sectioned by cryostat (Leica) into 20-μm coronal slices. Brain sections were stored in 0.01 M PBS (0.02% sodium azide) at 4°C before IHC.

For IHC, brain sections were blocked for 1 h at room temperature in 5% normal goat serum (NGS; Vector Laboratories, Burlingame, CA) diluted in 0.01 M PBS/0.1% Triton-X permeabilization buffer. Sections were then incubated for 48 h at 4°C in NGS blocking/permeabilization buffer, including primary antibodies raised against c-fos (rabbit anti-c-fos, Millipore, Danvers, MA, ABE457; 1:5000) and the NE transporter (NET; chicken anti-NET, #260006, Synaptic System, Goettingen, Germany; 1:3000). After washing in 0.01 M PBS, sections were incubated for 2 h in blocking/permeabilization buffer with goat anti-rabbit AlexaFluor 488 and goat anti-chicken AlexaFluor 568 (Invitrogen, Carlsbad, CA; 1:500). After washing, the sections were mounted onto Superfrost Plus slides and coverslipped with Fluoromount-G plus DAPI (Southern Biotech, Birmingham, AL).

### 2.5 Statistical analysis

The effects of the water vs predator odor on defensive burying and grooming in *Dbh +/-* and *Dbh -/-* mice were compared using repeated measures 2-way ANOVA (genotype x odorant), with post hoc Tukey tests for multiple comparisons where appropriate. The effects of drugs vs vehicle on time spent engaged in repetitive behaviors or locomotor activity were assessed using repeated measures 1-way ANOVA with post hoc Dunnett’s multiple comparison tests (SCH-23390) or paired sample t-tests (all other drugs). Contingency data of nestlet shredding and sleeping between genotypes were analyzed using Fisher’s exact tests. Although the current study was not sufficiently powered to detect sex differences, preliminary analysis of the data revealed roughly equivalent distributions and means between males and females (data not shown), similar to what we previously reported for behavioral and neurochemical responses to novel odors (Lustberg et al 2022). Thus, male and female data were pooled within each genotype and treatment group.

## 3. Results

### 3.1 Dbh -/- mice exhibit decreased defensive burying and increased grooming in response to predator odor

We first assessed the responses of *Dbh -/-* and NE-competent *Dbh +/-* control mice to predator odor and water by measuring defensive burying (**Fig. 1A, Supplemental movie 1**) and grooming (**Fig. 1B, Supplemental movie 1**). Control animals displayed modest levels of grooming during water exposure and no defensive burying. Predator odor dramatically increased defensive burying and total digging but had no impact on grooming in control mice. By contrast, *Dbh -/-* mice displayed high levels of grooming in response to water and were behaviorally indifferent to predator odor, with no emergence of defensive burying and no change in grooming.

**Fig. 1.**
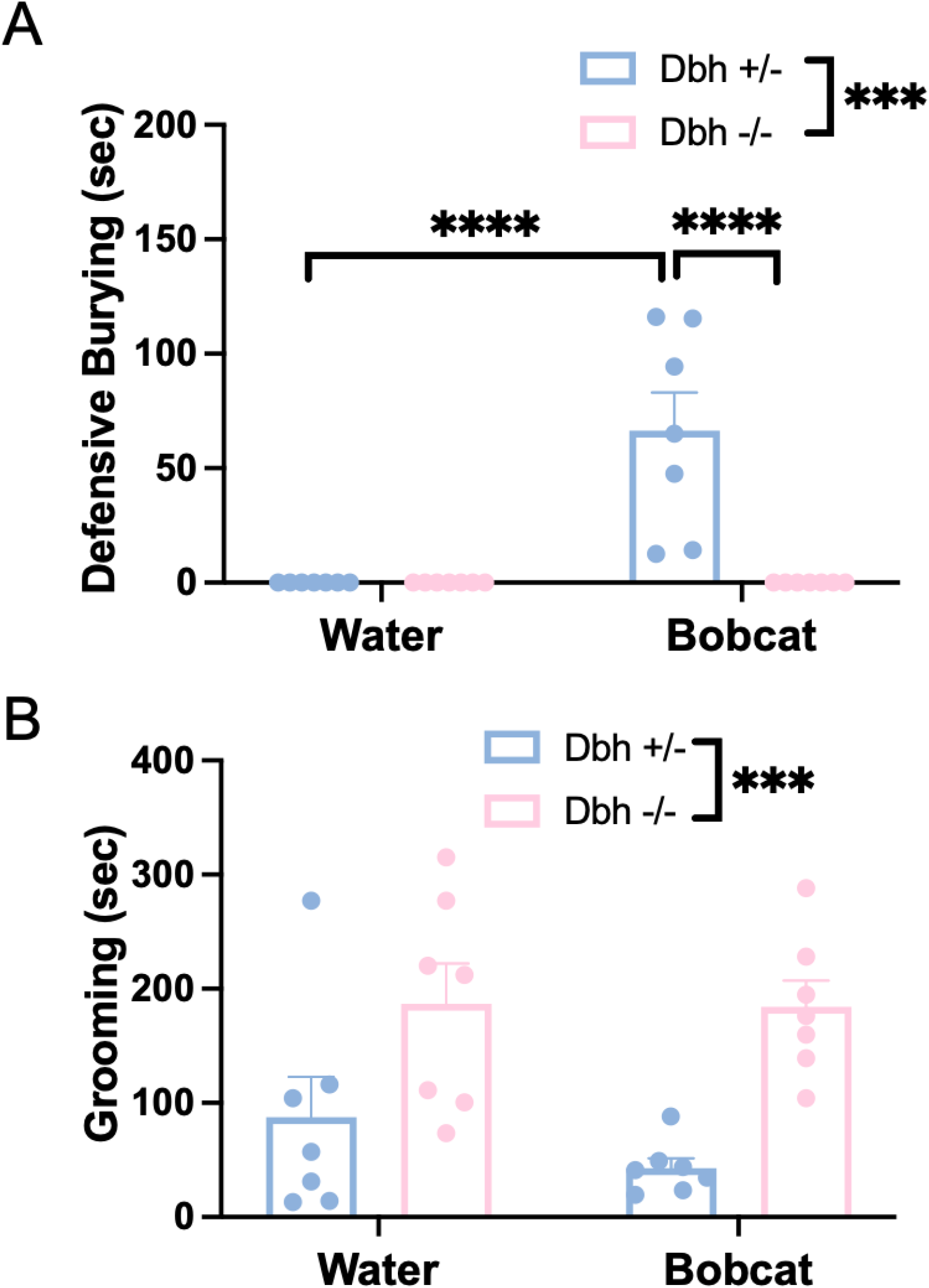
Effects of DBH knockout on behavioral responses to predator odor. *Dbh +/-* (n=7) and *Dbh -/-* (n=7) mice were exposed to water or bobcat urine for 10 min, and the amount of time engaged in defensive burying (**A**) and grooming (**B**) were assessed. Shown are mean ± SEM. ***p< 0.001, ****p<0.0001.

For defensive burying, there were main effects of genotype [F(1,24) = 15.96, p = 0.0005], odorant [F(1,24) = 15.96, p = 0.0005], and odorant x genotype interaction [F(1,24) = 15.96, p = 0.0005]. Post hoc analysis revealed that *Dbh +/-* mice showed significantly increased defensive burying (p < 0.0001) in the predator odor condition compared with water, and they also engaged in significantly more defensive burying (p < 0.0001) than the *Dbh -/-* mice in the predator odor condition. For grooming, there was a main effect of genotype [F(1,24) =18.82, p = 0.002], but no main effect of odorant [F(1,24) = 0.7293, p = 0.40] or odorant x genotype interaction [F(1,24) = 0.57, p = 0.46].

### 3.2 Adrenergic antagonists suppress predator odor-induced digging behavior in control animals

Our data from *Dbh -/-* mice suggested that NE is required for defensive burying in response to predator odor. To determine whether this is true in NE-competent mice as well as which receptors are involved, we next assessed the effects of specific AR antagonists on the predator odor responses of *Dbh +/-* mice. The compounds tested included prazosin (0.5 mg/kg, α1AR antagonist), atipamezole (1 mg/kg, α2AR antagonist), and propranolol (5 mg/kg, βAR antagonist). Compared with their respective vehicles, each compound tested significantly reduced burying (prazosin: t(7) = 2.46, p = 0.044; atipamezole: t(6) = 3.38, p = 0.015); propranolol: t(7) = 2.85, p = 0.025) (**Fig. 2A-C**), indicating that α1AR, α2AR, and βAR activation all contribute to this predator odor-induced defensive behavior. Prazosin was the only compound that attenuated grooming in control mice (**Fig. 2D-F**) (prazosin: t(7) = 3.03, p = 0.019; atipamezole: t(6) = 0.74, p = 0.49; propranolol: t(7) = 0.85, p = 0.43). Overall, these results indicated that defensive burying was the predator odor-induced behavior most sensitive to blocking NE transmission.

**Fig. 2.**
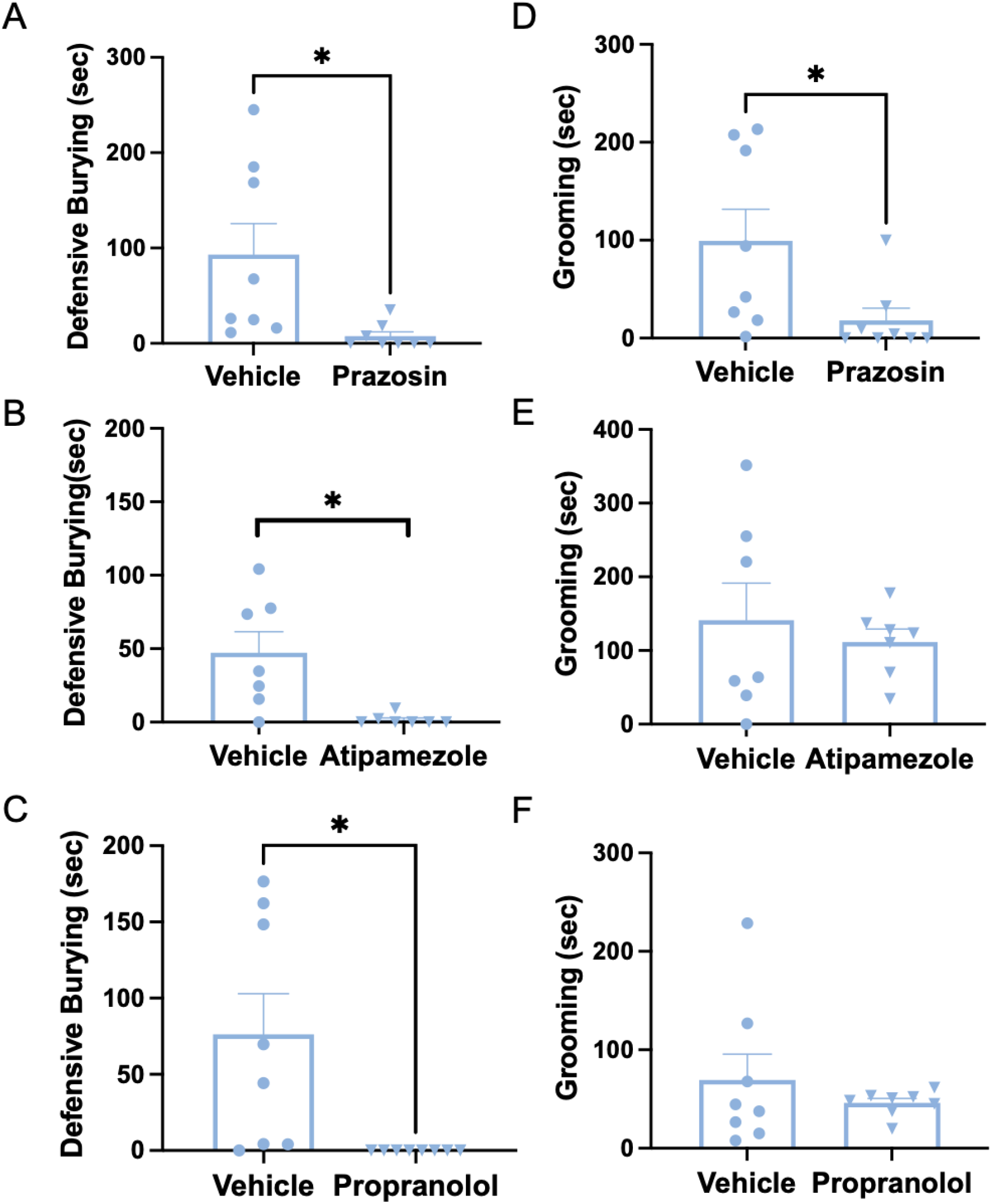
Effects of adrenergic drugs on behavioral responses to predator odor in control mice. *Dbh +/-* (n=8) mice were pretreated with vehicle, the α1AR antagonist prazosin (0.5 mg/kg), the α2AR antagonist atipamezole (1 mg/kg), or the βAR antagonist propranolol (5 mg/kg) and exposed to bobcat urine 30 min later for 10 min. Shown is the mean ± SEM time engaged in defensive burying (**A**-**C**) and grooming (**D**-**F**). *p<0.05.

### 3.3 Pharmacological DBH inhibition reduces defensive burying and increases grooming following exposure to predator odor in normal mice

Because AR antagonists recapitulated the decrease in digging behaviors but not the increase in grooming observed in the *Dbh -/-* mice, we reasoned that increased grooming was mediated by ectopic DA produced by LC neurons in the knockouts. To test this idea, we used the selective DBH inhibitor nepicastat, which unlike AR antagonists increases DA in addition to decreasing NE (Devoto et al 2015, Ohkubo et al 2019, Schroeder et al 2010, Stanley et al 1997). Nepicastat administration to *Dbh +/-* control mice phenocopied both aberrant responses of *Dbh-/-* mice to predator odor [defensive burying: t(9)=3.29, p = 0.009; grooming: t(9)=2.60, p = 0.029) (**Fig 3**), suggesting that increasing DA is necessary for abnormal *Dbh -/-* mouse grooming.

**Fig. 3.**
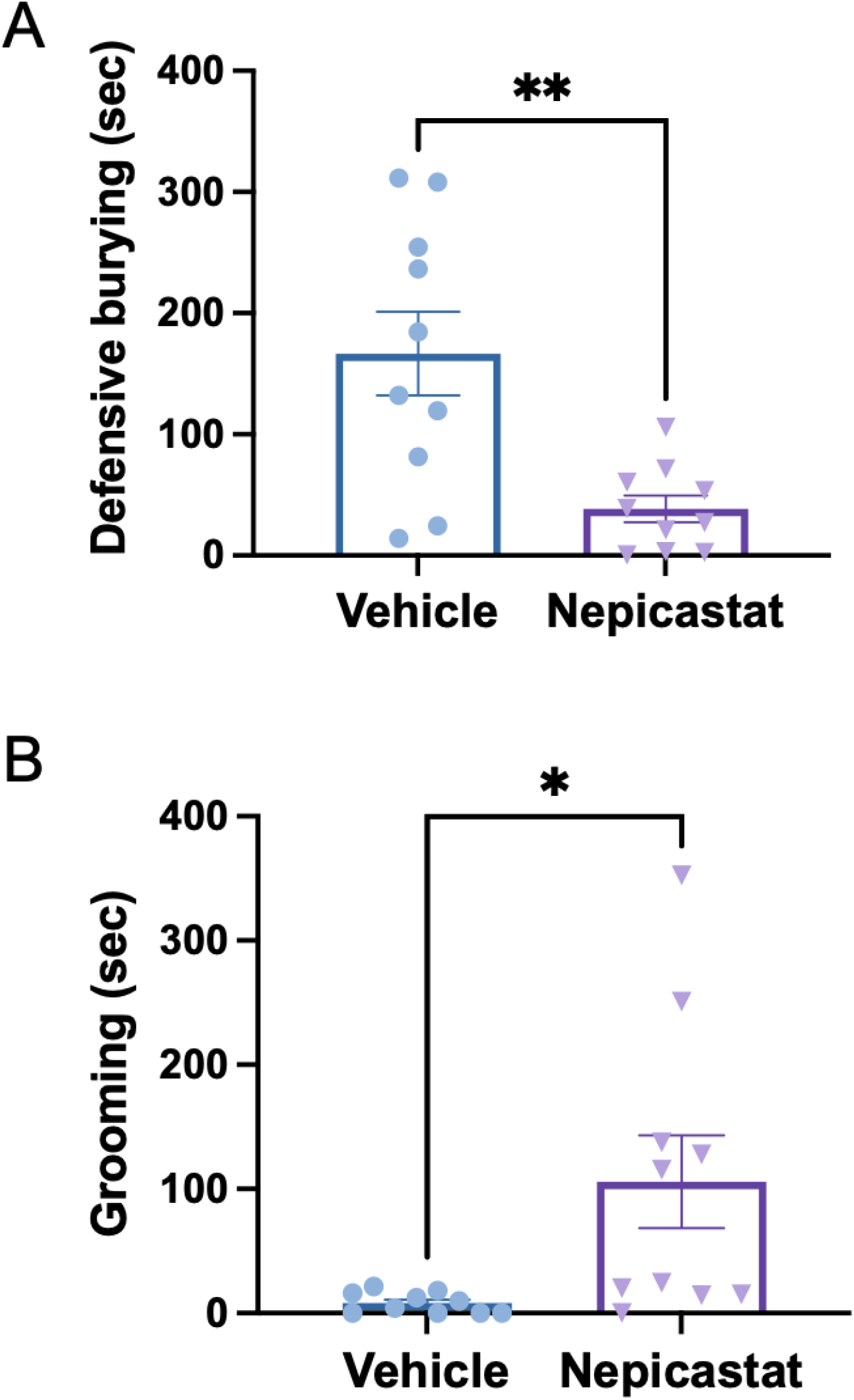
Effects of pharmacological DBH inhibition on behavioral responses to predator odor. *Dbh +/-* mice (n=10) were pretreated with vehicle or the DBH inhibitor nepicastat (100 mg/kg) and exposed to bobcat urine for 10 min, and the amount of time engaged in defensive burying (**A**) and grooming (**B**) were assessed. Shown are the mean ± SEM. *p< 0.05, **p<0.01.

### 3.4 DA receptor antagonists attenuate excessive grooming in Dbh -/- mice in the presence of predator odor

To directly test whether excessive DA transmission was required for grooming in *Dbh -/-* mice, we assessed the effects of DA receptor antagonists on predator odor response in the knockouts. The drugs tested included flupenthixol (0.25 mg/kg, nonselective DA receptor antagonist), L-741,626 (10 mg/kg, D2 receptor antagonist), and SCH-23390 (0.03 mg/kg, D1 receptor antagonist). Flupenthixol [t(12) = 2.76, p = 0.0037] (**Fig. 4A**) significantly reduced grooming in *Dbh -/-* mice in the presence of predator odor, while L-741,626 had no effect [t(7) = 0.75, p = 0.48] (**Fig. 4B**). SCH-23390 completely eliminated grooming, but we also noted that mice administered SCH-23390 ceased nearly all movement. SCH-23390-treated mice were capable of moving and did so when gently prodded, but failed to ambulate volitionally. In an attempt to separate the effects of D1 blockade on grooming and general motor activity, we also tested two lower doses (0.01 and 0.006 mg/kg), and the results are shown in **Fig. 4C** and **4D**. There was a main effect of dose on both grooming [F(1.701, 11.90) = 40.42, p < 0.0001], and locomotor activity [F(1.226, 8.579) = 15.59, p = 0.0028]. Post hoc analysis revealed that the 0.03 dose fully suppressed grooming [q(7) = 12.66, p < 0.0001] and locomotor activity [q(7) = 8.48, p = 0.002]. Similarly, there was almost no grooming following administration of the 0.01 mg/kg dose [q(7) = 8.83, p = 0.0001], while locomotor activity returned to ∼50% of saline control [q(7) = 7.06, p = 0.005]. Finally, the 0.006 mg/kg dose induced a borderline significant reduction of grooming [q(7) = 2.81, p = 0.06] with no effect on locomotor activity [q(7) = 0.06, p = 0.99]. These results suggest that excessive grooming in *Dbh -/-* mice is mediated by D1, but not D2 transmission, although we cannot completely rule out a non-specific effect. Vehicle-treated *Dbh -/-* mice exhibited almost no defensive burying (see **Fig. 1A**), and none of the dopaminergic antagonists altered this outcome (data not shown).

**Fig. 4.**
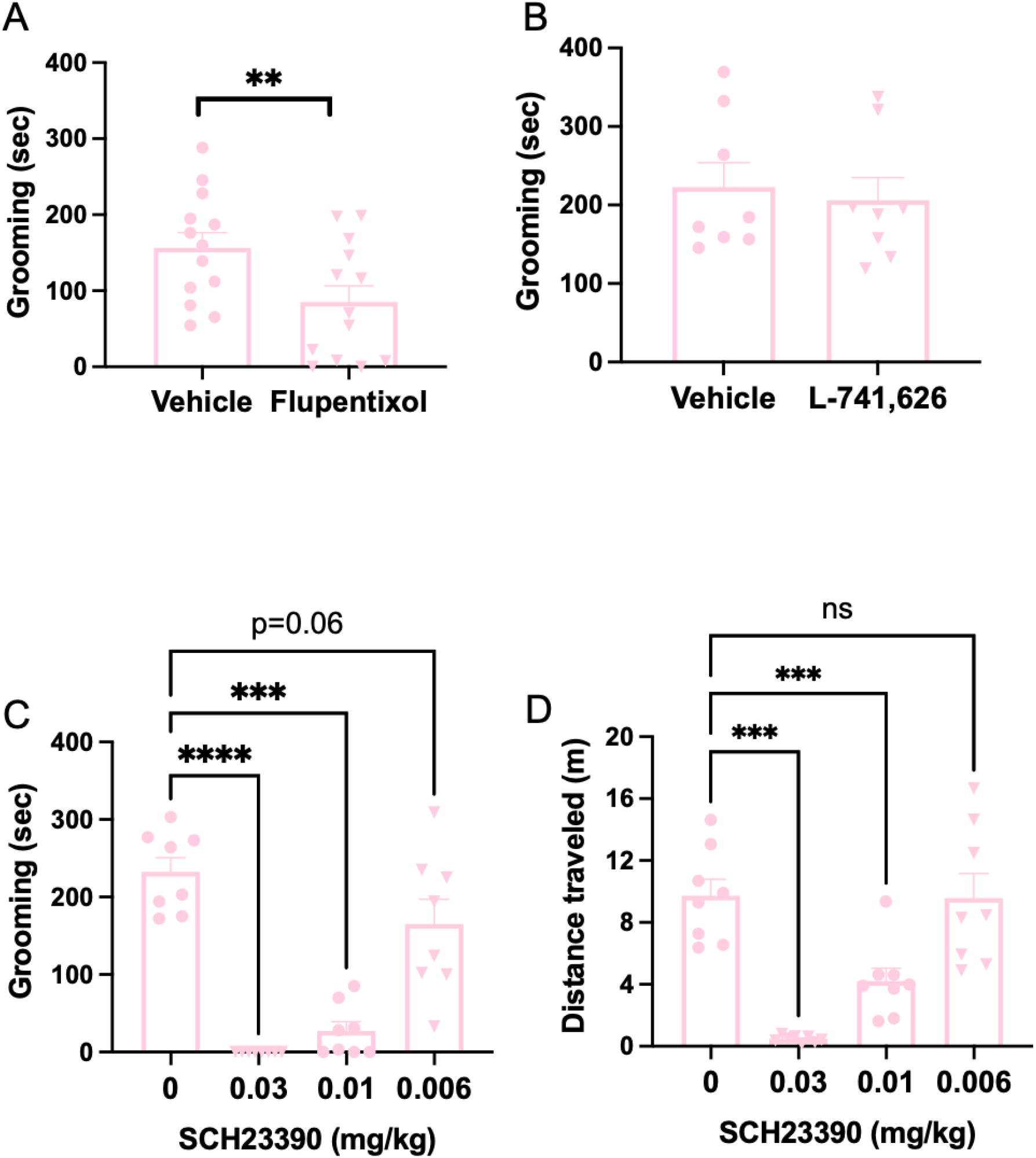
Effects of dopaminergic drugs on predator odor-induced grooming and locomotor activity in *Dbh -/-* mice. *Dbh -/-* mice were pretreated with vehicle (n=7-13), the non-selective DA receptor antagonist flupenthixol (0.25 mg/kg; n=7), the D2 antagonist L-741,626 (10 mg/kg; n=8), or the D1 antagonist SCH-23390 (0.03, 0.01, or 0.006 mg/kg; n=8), or and exposed to bobcat urine 30 min later for 10 min. Shown is the mean ± SEM time engaged in grooming (**A-C**) and distance travelled (**D**). **** p<0.0001, ***p<0.001, **p<0.01, ns=not significant.

### 3.5 α1AR blockade has no effect on behavioral response to predator odor in Dbh -/- mice

Our data thus far have indicated that NE transmission was responsible for defensive burying, while DA promoted grooming. However, because we observed a significant reduction in grooming in the *Dbh +/-* mice treated with prazosin, and DA can signal through α1ARs under some conditions (Cilz et al 2014, Ozkan et al 2017, Paladini et al 2001), we speculated that some of the excessive grooming observed in *Dbh -/-* mice might be driven by DA-mediated α1AR transmission. To test this hypothesis, we administered prazosin (0.5 mg/kg) to *Dbh -/-* mice in the presence of predator odor, but failed to observe a drug effect on grooming (vehicle: 223.2 ± 49.85 sec; prazosin 216.3 ± 45.56 sec; p = 0.8986). Combined with the outcomes from the DA receptor antagonist experiments, these results suggest that while α1ARs modestly contribute to predator odor-induced grooming in normal mice, they are not required for the excessive grooming response of *Dbh -/-* mice, which appears to be mediated by excessive activation of D1 receptors.

### 3.6 Dbh -/- mice lack predator odor-induced arousal and aversion

Predator odors typically elicit innate vigilance and aversion in prey that is resistant to habituation. We and others have shown that, when exposed to a novel environment and given access to a cotton nestlet, normal mice rapidly shred the nestlet, build a nest, and fall asleep in it (Lustberg et al 2020a). To examine the effects of disrupting LC catecholamine homeostasis on these responses to predator odor, we measured nestlet shredding and arousal state in *Dbh +/-* and *Dbh -/-* mice after 90 min of exposure to a novel environment with a nestlet treated with bobcat urine. To control for the influence of a novel odor, a separate cohort of mice experienced the same paradigm except that a non-threatening but unfamiliar odor (lavender oil) was used instead of bobcat urine. In the presence of lavender, most (6/8) *Dbh +/-* mice shredded the nestlet (**Fig. 5A**) and were asleep (**Fig. 5B**) in the nest they built by the 90 min time point. As we have shown previously with unscented nestlet (Lustberg et al 2020a), *Dbh -/-* mice failed to shred the nestlet (0/7) (**Fig. 5A**) but still fell asleep (6/7) (**Fig. 5B**). Fisher’s exact tests revealed a significant difference between genotypes for shredding (p=0.007) but not sleeping (p>0.99).

**Fig. 5.**
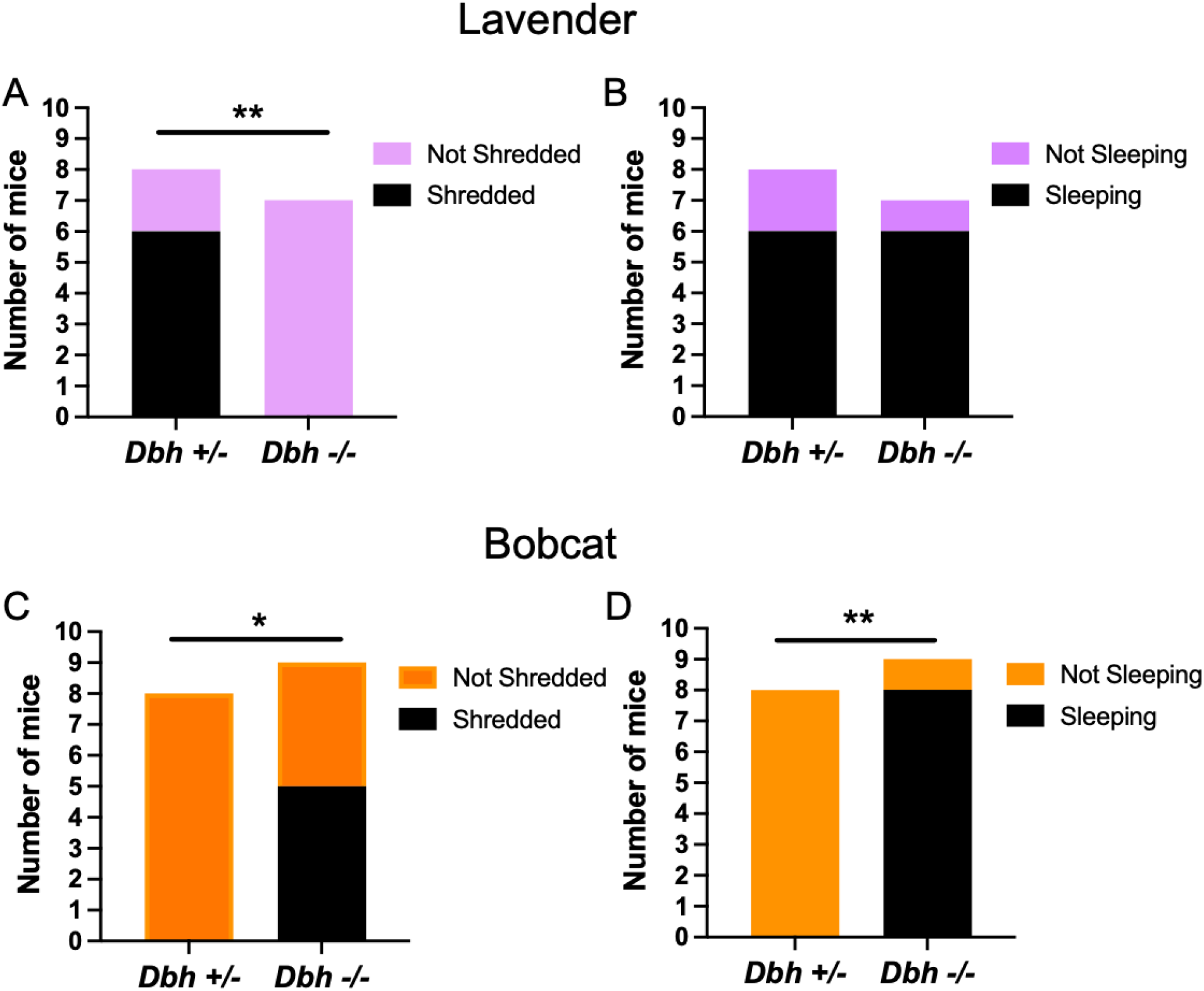
Effects of DBH knockout on arousal and nestlet shredding during predator odor exposure. *Dbh +/-* and *Dbh -/-* mice were exposed to lavender oil (**A**-**B;** n=7,8) or bobcat urine (**C**-**D**; n=8,9) for 90 min, and the fraction of animals that shredded their cotton nesting material (**A**,**C**) and fell asleep (**B**,**D**) were assessed. *p< 0.05, **p<0.01.

In contrast to lavender odor, none (0/8) of the *Dbh +/-* mice shredded their nestlet (**Fig. 5C**) or fell asleep (**Fig. 5D**) in the presence of bobcat urine. The behavior of *Dbh -/-* mice also changed when exposed to predator odor, but in unexpected directions. While *Dbh -/-* mice never engaged in shredding unscented or lavender-scented nestlets within 90 min, a majority (5/9) of them fully (100%) shredded the bobcat urine-soaked nestlet (**Fig. 5C**). Those that did build a nest fell asleep in it, and altogether 8/9 of the *Dbh -/-* mice were asleep by the end of the test (**Fig. 5D**). Fisher’s exact tests revealed significant differences between genotypes for shredding (p=0.0294) and sleeping (p=0.0004). Pilot experiments using different source of predator odor (fox urine and tufts of domestic cat fur instead of bobcat urine) also revealed paradoxical appetitive responses in knockout mice; *Dbh +/-* animals engaged in typical defensive burying behavior, while most *Dbh -/-* mice instead sat on top of the cat fur and groomed themselves (**Supplemental movie 2**). These results demonstrate that predator odor elicits powerful arousal and aversion in normal mice, and that loss of DBH activity produces aberrant behavioral responses that include apparent affiliative properties.

### 3.8 c-fos immunohistochemistry reveals predator odor-responsive brain regions that differ between Dbh +/- and Dbh -/- mice

To identify candidate brain regions that are activated by predator odor and facilitate differential responses in *Dbh +/-* and *Dbh -/-* mice, c-fos immunohistochemistry was conducted in animals exposed to a novel environment with a bobcat urine-treated nestlet. Analysis focused on the LC and other brain regions that receive innervation from noradrenergic neurons and have been implicated in predator odor responses, including the anterior cingulate cortex (ACC), dorsal BNST (dBNST), periaqueductal gray (PAG), periventricular nucleus of the hypothalamus (PVN), lateral septum (LS), and medial amygdala (MeA) (**Fig. 6**). Repeated measures 2-way ANOVA showed a main effect of brain region [F(3,45) = 76.57, p < 0.0001] and a genotype x brain region interaction [F(6,90) = 44.84, p < 0.0001]. Sidak’s post hoc tests revealed that *Dbh +/-* mice had significantly more c-fos+ cells than *Dbh -/-* mice in the ACC (t = 8.361, p < 0001), dBNST (t = 7.767, p < 0.0001), PAG (t = 9.897, p < 0.0001), and LS (t = 7.460, p < 0.001), while

**Fig. 6.**
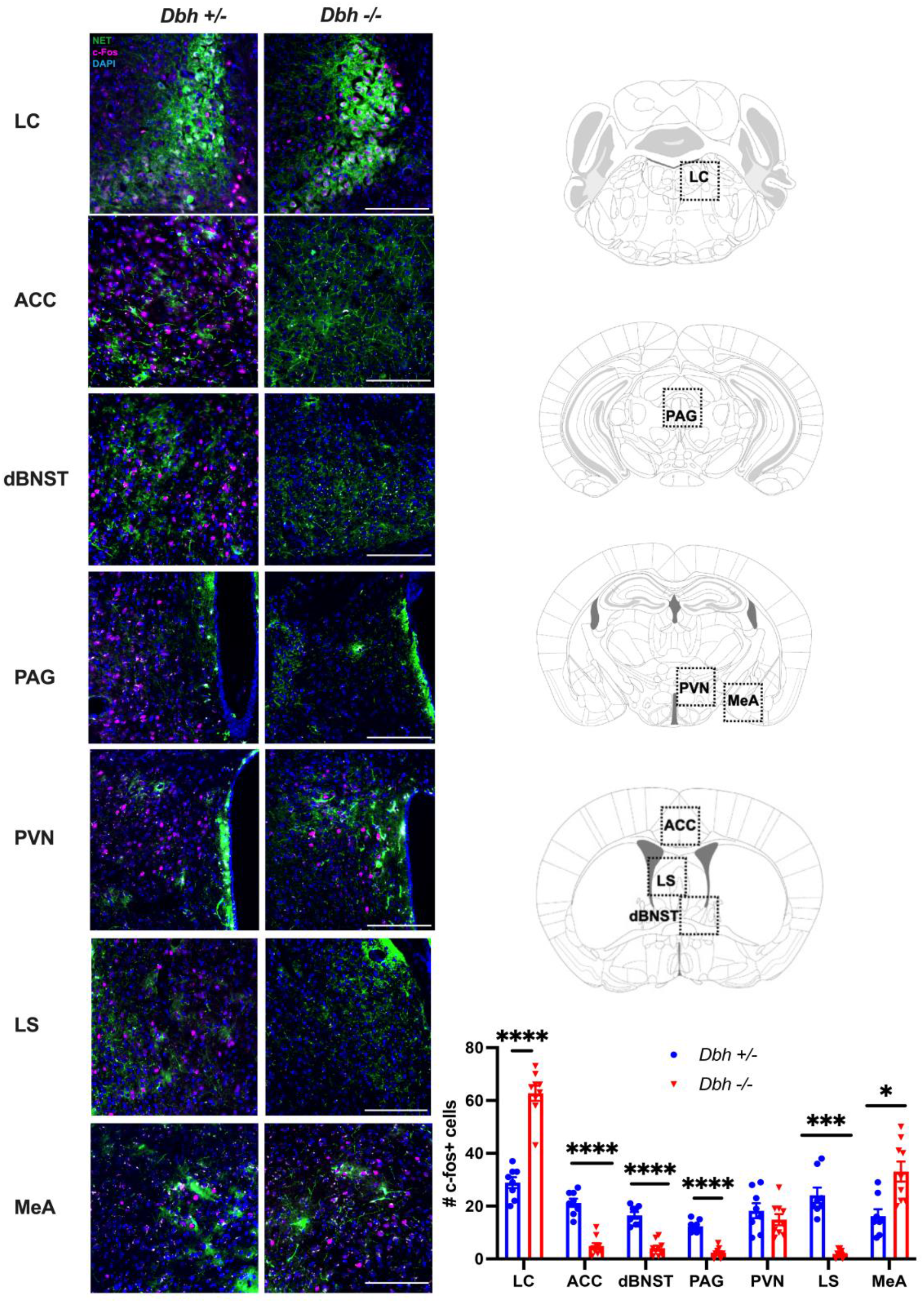
Regional c-fos responses to predator odor in *Dbh +/-* and *Dbh -/-* mice. *Dbh +/-* (n=8) and *Dbh -/-* (n=9) mice were exposed to bobcat urine for 90 min. Shown are (A) representative images and (B) the mean ± SEM number of c-fos+ (magenta) cells in the locus coeruleus (LC), anterior cingulate cortex (ACC), dorsal bed nucleus of the stria terminalis (dBNST), periaqueductal gray (PAG), paraventricular nucleus of the hypothalamus (PVN), lateral septum (LS), and medial amygdala (MeA). Images are counterstained with the nuclear marker DAPI (blue) and the noradrenergic terminal marker norepinephrine transporter (NET; green). *p<0.05, ***p<0.001, ****p<0.0001.

*Dbh -/-* mice had significantly more c-fos+ cells in the LC (t = 9.554, p < 0.0001) and MeA (t = 3.679, p = 0.0177). No genotype differences were found in the PVN (t = 0.956, p = 0.9542). Integrated with the effects of genotype and drugs, these results imply that noradrenergic transmission in the ACC, dBNST, PAG, and/or the LS contribute to aversive/defensive effects of predator odor, while D1 signaling in the MeA promotes grooming and appetitive responses.

## 4. Discussion

### 4.1 Summary

Predator odors (or components thereof) such as bobcat urine and TMT are ethologically relevant stressors that elicit intense, innate behavioral responses in mice (Ferrero et al 2011, Janitzky et al 2015). In this study, we investigated the effects of manipulating catecholamine transmission on predator odor-induced stress in mice by comparing *Dbh -/-* mice, which lack NE and instead produce DA in their “noradrenergic” neurons, to their *Dbh +/-* littermates with normal catecholamine content. We also used pharmacological manipulation to target NE and DA signaling, allowing us to assess the effects of both chronic and acute catecholamine manipulation.

### 4.2 Adrenergic control of predator odor-induced defensive burying

Defensive burying is considered an active coping strategy in response to stress, in which the mouse attempts to remove or avoid aversive stimuli. In the shock probe defensive burying test, mice will use bedding to bury electrified prods and remove the source of shock pain (Tillage et al 2020). The marble burying test has also demonstrated that burying is stress-sensitive, with stressed mice burying more marbles than unstressed mice (Kedia & Chattarji 2014), and we have shown that *Dbh -/-* mice bury fewer marbles than controls (Lustberg et al 2020a).

We found that both *Dbh -/-* and *Dbh +/-* mice engage in similarly low levels of defensive burying when exposed to a novel cage with a water-soaked cotton nestlet, indicating that this environment is not particularly stressful. *Dbh +/-* mice show increased defensive burying when exposed to predator odor as compared to water, suggesting that this behavior is an innate adaptive defensive response to threat. The combined results from our experiments with *Dbh -/-* mice, acute pharmacological DBH inhibition, and adrenergic receptor antagonists demonstrate that noradrenergic transmission is required for defensive burying during predator odor exposure. We previously showed that NE is critical for digging in the presence of novel odors but did not precisely identify the receptor subtypes responsible (Lustberg et al 2022).

The effects of βAR and α1AR activation are partially overlapping but can also be distinct. βAR activation is necessary for many types of anxiety-like and defensive behaviors, including responses to physical threats (e.g. shock) (Bondi et al 2007), psychological threats (e.g. novel environments) (Lustberg et al 2022), and anxiogenic drugs (e.g. cocaine) (Schank et al 2008), while α1AR signaling promotes both exploratory and aversive behaviors in novel environments (Lustberg et al 2020a, Stone et al 2006, Stone et al 1999). α1AR blockade decreases defensive behavior in the presence of a traumatic cue in rats (Ketenci et al 2020), and antagonism of βARs attenuates cocaine withdrawal-induced defensive burying in rats (Harris and Aston-Jones, 1993). Interestingly, the βAR antagonist propranolol blocks the anxiogenic effects of optogenetic LC stimulation in the elevated zero maze, while only the α1AR antagonist prazosin prevents real-time place aversion (McCall et al 2015). We have shown that propranolol diminishes nestlet shredding, while prazosin attenuates marble burying in a novel environment (Lustberg et al 2020a). By contrast, antagonism of either receptor reduces defensive burying in the shock probe test. Thus, the current finding that both βARs and α1ARs contribute to defensive burying behavior during predator odor exposure is consistent with previous literature.

The ability of the α2AR antagonist atipamezole to reduce predator odor-induced defensive burying was more of a surprise. α2ARs are expressed widely throughout the brain but are enriched in LC neurons, where they function as inhibitory autoreceptors (Aghajanian & VanderMaelen 1982, Timmermans & van Zwieten 1982). Thus, α2AR agonists often decrease, while α2AR antagonists can increase NE transmission and anxiety-like behaviors (Lustberg et al 2020a, Lustberg et al 2020b). In the current study, atipamezole had effects similar to those of propranolol and prazosin, suggesting that it was not primarily acting on α2AR autoreceptors but rather by blocking α2AR heteroreceptors on LC target cells. α2ARs are expressed in stress-sensitive brain regions such as the prefrontal cortex (PFC, which includes the ACC) and the dBNST, where they can have paradoxical excitatory effects on neurons and networks (Harris et al 2018, Wang et al 2007). For example, stimulation of postsynaptic α2ARs in the dBNST enhances neuronal activity and promotes anxiety-like behavior in the elevated plus maze (Harris et al 2018). We speculate that atimpamezole-induced reduction of defensive burying in response to predator odor is mediated by similar mechanisms.

### 4.3 Dopaminergic control of grooming

Grooming, unlike digging, is not typically considered a stress coping response, as mice will groom in situations of both comfort and distress (Smolinsky AN 2009). Indeed, cat odor often inhibits grooming in rodents (McGregor et al 2004). Repetitive, purposeless self-grooming is often observed in animal models of obsessive-compulsive disorder (OCD), Tourette’s syndrome (TS), and autism spectrum disorder (ASD) (Lewis 2011, Nordstrom & Burton 2002, Welch et al 2007). DA signaling has been previously implicated in facilitating grooming behavior. For example, D1, but not D2, receptor signaling contributes to stereotyped self-grooming in rats (Berridge & Aldridge 2000a, Berridge & Aldridge 2000b) and novelty-induced grooming in mice (Drago et al 1999).

As a precursor for NE, DA is produced in noradrenergic neurons. Due to incomplete conversion to NE by DBH, DA can also be released by these cells and mediates some LC functions including specific forms of learning and memory (Chowdhury et al 2022, Kempadoo et al 2016, Wagatsuma et al 2018). *Dbh -/-* mice lack NE and instead exclusively release DA from LC neurons under conditions that normally result in NE transmission. Converging lines of evidence from our study indicate that the excessive grooming displayed by *Dbh -/-* mice is mediated by DA. First, high levels of grooming in the presence of predator odor were observed in control mice treated with the DBH inhibitor nepicastat, but not in mice following AR blockade. Second, grooming behavior in *Dbh -/-* mice was attenuated by DA receptor antagonists.

Our findings that the non-selective DA receptor antagonist flupenthixol and the D1-selective antagonist SCH-23390, but not the selective D2 antagonist L-741,626 blocked grooming in *Dbh -/-* mice exposed to predator odor indicates this behavior is mediated by D1 receptors. One caveat to this interpretation is that D1 blockade is well known to suppress many aspects of motor activity, including cocaine-induced hyperactivity (Cabib et al 1991) and rearing in a novel cage (Dreher & Jackson 1989). Another study using SCH-23390 also showed that it can induce catalepsy (Chinen & Frussa-Filho 1999), although at a much higher dose (0.1 mg/kg) than the one we used. In our experiments, the effects of SCH-23390 on grooming were more potent than those on locomotion, but there was no dose that significantly reduced grooming without effects on distance travelled; the closest was 0.006 mg/kg, which allowed normal locomotion while almost reaching significance for grooming (p = 0.06). Prazosin reduced grooming in *Dbh +/-* but not *Dbh -/-* mice, indicating that NE signaling through α1ARs promote this behavior exclusively in normal animals. This is consistent with our previous finding that prazosin reduces amphetamine-induced locomotion in *Dbh +/-* but not *Dbh -/-* mice (Weinshenker et al 2002), and further implicates D1 receptor mediation in knockouts.

We previously showed that *Dbh -/-* mice groom more than *Dbh +/-* mice in the presence of a novel but non-threatening odorant, and this excessive grooming was reduced by administration of flupentixol (Lustberg et al 2022). Therefore, the results from our genetic and pharmacological experiments suggest that the neurochemistry and circuity that drive grooming in the presence of predator odor resemble those controlling grooming responses to novel odors and/or environments.

### 4.4 Dbh -/- mice have impaired arousal and aversion responses to predator odor

Both NE and DA promote arousal. Activation of noradrenergic neurons in the LC and dopaminergic neurons in the ventral tegmental area (VTA) and ventral PAG (vPAG) induce wakefulness, while their suppression reduces arousal (Carter et al 2010, Cho et al 2017, Eban-Rothschild et al 2016, Lu et al 2006, Poe et al 2020, Porter-Stransky et al 2019). Many environmental challenges such as low dose amphetamine, white noise, and human handling that increase wakefulness in normal mice fail to do so in *Dbh -/-* mice (Hunsley & Palmiter 2003, Hunsley & Palmiter 2004), CRISPR-mediated knockout of DBH in the mouse LC attenuates arousal triggered by rat bedding (Yamaguchi et al 2018), and AR antagonists can also reduce wakefulness in control mice (Mitchell & Weinshenker 2010, Porter-Stransky et al 2019).

Sustained arousal is an adaptive response to threats such as predator odor. Normal mice exposed to a new cage with an unscented cotton nestlet will initially explore the novel environment, then shred the material and fall asleep in the nest within 60-90 min (Lustberg et al 2020a). In the current study, we showed that while most control mice shredded a cotton square containing a novel but non-threatening odor (lavender) and fell asleep in the nesting material, none shredded a bobcat urine-soaked nestlet and all remained vigilant for at least 90 min. *Dbh -/-* mice had a paradoxical response to novel and threatening odors. The knockouts fail to shred odorless (Lustberg et al 2020a) or lavender-scented (this study) material, but remarkably built a nest and fell asleep in it only when it contained bobcat urine. Because defensive burying and grooming responses to predator odor in *Dbh -/-* mice did not differ from those to plain water, it was not initially clear whether *Dbh -/-* mice could even detect bobcat urine, However, the nestlet result refutes that possibility, and odor sensation in general is intact in both *Dbh -/-* mice and people with complete DBH deficiency (Garland et al 2011, Lustberg et al 2022, Thomas & Palmiter 1997). Therefore, our findings suggest that suppression of DBH produces aberrant responses of rodents to threats. Crucially, NE transmission is generally anxiogenic and mediates aversive responses to stress, whereas DA signaling is a key component of many appetitive/approach behaviors (Iordanova et al 2021). Thus, we speculate that the loss of NE suppresses adaptive fearful/defensive behaviors, while the increased DA reverses the valence of predator odor from aversive to appetitive. Further experiments combing genetic and pharmacological manipulations of catecholamine signaling and circuits with real-time and conditioned place preference/aversion approaches will help test this idea.

### 4.5 Neuroanatomical substrates of NE and DA action

Results from the c-fos immunostaining experiment provide clues about where NE and DA released from noradrenergic neurons may drive predator odor-induced behaviors (**Fig. 7**). *Dbh -/-* mice had fewer c-fos+ neurons in a field of NET+ fibers in the ACC, dBNST, PAG, and LS, suggesting that NE is required in one or more of these brain regions for predator odor-induced defensive behaviors (i.e. burying, arousal). These regions were chosen for examination because they are activated by predator odor, are innervated by noradrenergic neurons, and/or are key nodes in defensive behavior circuits (Janitzky et al 2015, Jhang et al 2018, McGregor et al 2004). Because the contribution of these regions and circuits to predator odor responses is vast and has been reviewed elsewhere (Dielenberg & McGregor 2001, Rosen et al 2008, Takahashi 2014), we will only highlight a few relevant findings with particular relevance to our data here. For example, microinfusion of either βAR or α1AR antagonists into the LS reduced defensive burying in the shock probe test, while intra-BNST application of the α2AR agonist clonidine suppressed TMT-induced NE release and fear responses (Fendt et al 2005).

**Fig. 7.**
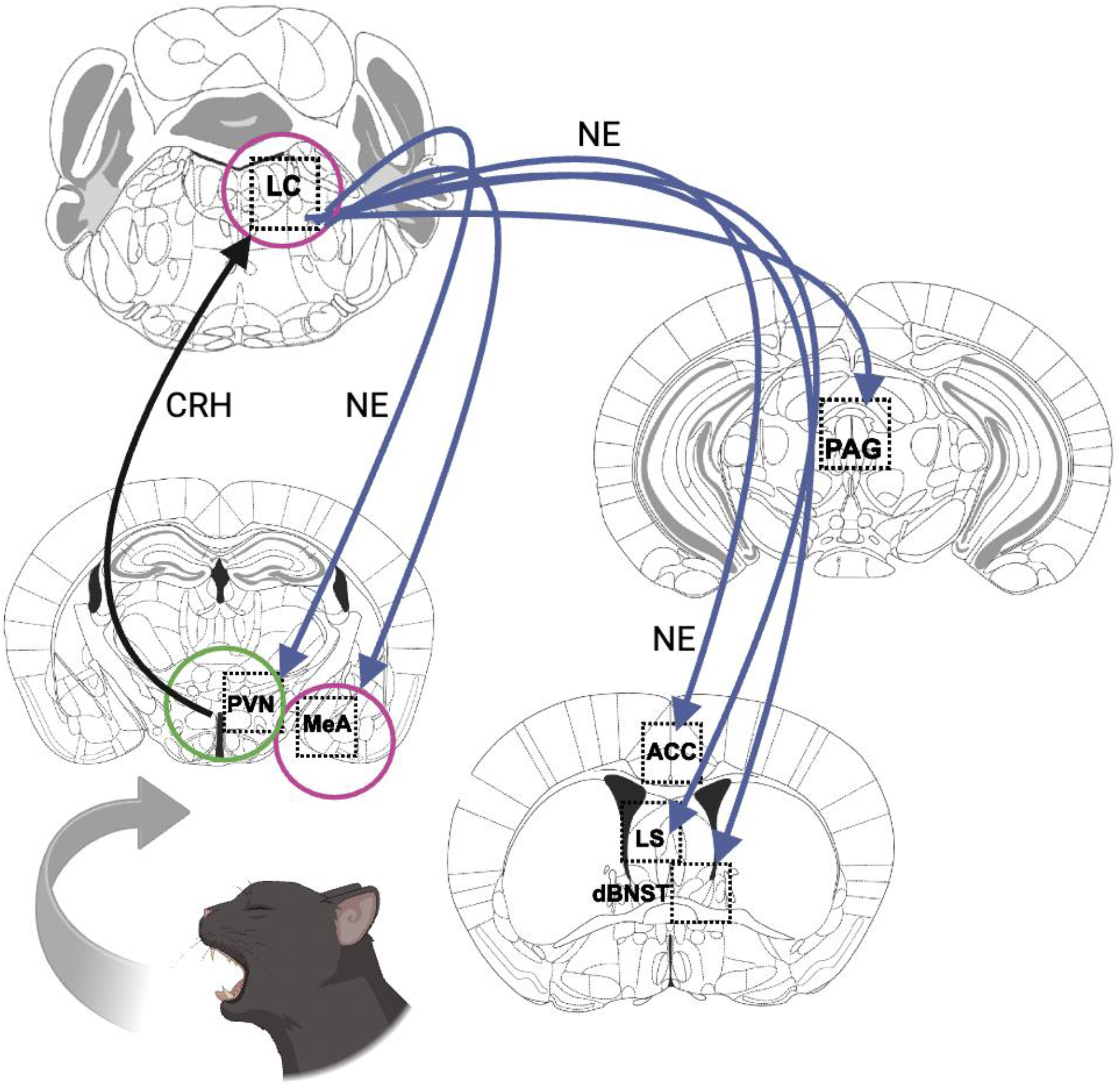
Proposed noradrenergic circuit for defensive behavioral responses to predator odors. The PVN hypothalamus is activated by ancient odorant pathways, in turn exciting the LC by releasing CRH (black arrow). The LC projects to a constellation of brain regions implicated in defensive behavior, where it releases NE in DBH-sufficient animals (blue arrows). Loss of NE results in hypoactivity in PAG, ACC, LS, and dBNST (regions without circles). Interestingly, loss of NE did not affect neuronal activity in the PVN (green circle), suggesting this pathway remains intact. Further, NE deficiency resulted in hyperactivity in the LC and the MeA (magenta circles). Image created with Biorender.

Furthermore, BNST inactivation not only suppresses aversive responses to cat urine in rats but increases duration of contact with the odor source, suggesting enhanced affiliative behavior similar to those observed in *Dbh -/-* mice (Xu et al 2012). The PAG is generally implicated in defensive behaviors and fear responses and is of particular interest here because we previously identified a noradrenergic LC-PAG circuit regulating arousal that may be engaged to maintain vigilance under threatening conditions (Porter-Stransky et al 2019). The lack of NE in *Dbh -/-* mice impairs this circuit and leads to reduced arousal, which could explain why the knockouts fall asleep in the presence of bobcat urine. Interestingly, inactivation of the ACC increases predator odor-induced freezing, while activation has the opposite effect (Jhang et al 2018). It is possible that the increased activation of the ACC in our control mice was permissive for active coping (defensive burying) at the expense of passive coping (freezing). We did not observe a high degree of freezing in *Dbh -/-* mice despite the reduced c-fos response, suggesting that other brain regions responsible for grooming dominated in the knockouts.

Two regions, the MeA and LC, had an increased c-fos response in *Dbh -/-* mice. The MeA receives direct and indirect input from the olfactory system, as well as DA neurons in the VTA. Dopaminergic signaling within the MeA mediates social reward and has even been implicated in human bonding (Atzil et al 2017, Hu et al 2021, Takahashi 2014). D1 receptor-expressing MeA neurons are activated by cat urine, and facilitating VTA dopaminergic transmission or activating D1 MeA GABAergic outputs to the BNST causes animals to display increased approach to a predator odor threat (McGregor et al 2004, Miller et al 2019, Vincenz et al 2017).

These data are consistent with the idea that bobcat urine elicits paradoxical appetitive rather than aversive responses in *Dbh -/-* mice because of ectopic DA release from LC afferents in the MeA. Because α2AR inhibitory autoreceptors limit LC activation, it is likely that the increased c-fos response in the LC of *Dbh -/-* mice is due to their lack of NE and α2AR engagement (Paladini et al 2007).

### 4.6 Translational significance and implications for T. gondii infection

It is important to identify the neurobiological underpinnings of species-typical and maladaptive responses to threats because dysregulated behavioral responses to stressful stimuli represent many hallmark features of neuropsychiatric disorders, including posttraumatic stress disorder (PTSD), anxiety, and drug abuse (Dias et al 2013). Indeed, drugs that impact catecholaminergic transmission are used to treat affective diseases, although the mechanisms of action are not fully understood.

The results of this study may also have implications for *Toxoplasma gondii* infection. As a parasite that reproduces in cats as part of its life cycle, *T. gondii* impairs the innate fear of cat odor in rodents, increasing the chance that the parasite reaches its intermediate (feline) and permanent (larger animals, including humans) host (Kannan et al 2010, Vyas et al 2007, Webster et al 1994). Although the mechanisms by which *T. gondii* changes host behavior are not fully understood, recent studies showed that *T. gondii* infection in rodents profoundly decreases expression of DBH, reducing NE and elevating DA in the brain (Alsaady et al 2019,

Tedford et al 2023). Our results provide additional evidence that the *T. gondii*-induced reduction of DBH may lead to suppression of defensive behaviors that would normally promote escape from a predator. Indeed, the apparent affiliation of *Dbh -/-* mice to feline scents (i.e. sitting on domestic cat fur, nesting in bobcat urine-soaked cotton) may actually attract them to danger and would make them easy prey in the wild. We predict that at least some of the behavioral effects of *T. gondii* infection would be abolished or occluded in Dbh -/- mice. Since NE has anti-inflammatory properties (Chalermpalanupap et al 2013, Feinstein et al 2016), it is possible that *T. gondii*-induced inflammation would be greater in *Dbh -/-* mice. Consistent with our intriguing finding of increased c-fos in the MeA of *Dbh -/-* mice, reversal of *T. gondii-*induced epigenetic and gene expression signatures in this structure normalized defensive behavior and abolished the gain of attraction to cat odor (Hari Dass & Vyas 2014), further indicating a potential neuroanatomical substrate for the excessive DA in *Dbh -/-* mice.

*T. gondii*-induced changes in behavioral responses to predator stress is not merely a rodent curiosity; approximately one-third of the human population carries a chronic *T. gondii* infection, and while most infected people are grossly asymptomatic, toxoplasmosis can have devastating effects on the fetal brain and is associated with greater risk for schizophrenia, OCD, ASD, suicide, and homicide in people with latent *T. gondii* infection (Contopoulos-Ioannidis et al 2022, Cook et al 2015, Nayeri et al 2022, Nayeri et al 2020, Torrey & Yolken 2003, Virus et al 2021). Thus, further study of NE and DA circuitry underlying *T. gondii*-induced behavioral changes in rodents and of catecholaminergic drugs as therapies for disorders with stress-responsive repetitive behaviors are warranted.

## Acknowledgements

We thank Q. Eastman for helpful editing of the manuscript and Cordelia and Lila for donating fur. This work was supported by the National Institutes of Health (DA049257, AG061175, and AG079199 to DW; NS007480-20 to DL).

## Bibliography

Aghajanian GK, VanderMaelen CP. 1982. alpha 2-adrenoceptor-mediated hyperpolarization of locus coeruleus neurons: intracellular studies in vivo. Science 215: 1394–6

Alsaady I, Tedford E, Alsaad M, Bristow G, Kohli S, et al. 2019. Downregulation of the Central Noradrenergic System by Toxoplasma gondii Infection. Infect Immun 87

Atzil S, Touroutoglou A, Rudy T, Salcedo S, Feldman R, et al. 2017. Dopamine in the medial amygdala network mediates human bonding. Proc Natl Acad Sci U S A 114: 2361–66

Beas BS, Wright BJ, Skirzewski M, Leng Y, Hyun JH, et al. 2018. The locus coeruleus drives disinhibition in the midline thalamus via a dopaminergic mechanism. Nat Neurosci 21: 963–73

Berridge KC, Aldridge JW. 2000a. Super-stereotypy I: enhancement of a complex movement sequence by systemic dopamine D1 agonists. Synapse 37: 194–204

Berridge KC, Aldridge JW. 2000b. Super-stereotypy II: enhancement of a complex movement sequence by intraventricular dopamine D1 agonists. Synapse 37: 205–15

Bondi CO, Barrera G, Lapiz MD, Bedard T, Mahan A, Morilak DA. 2007. Noradrenergic facilitation of shock-probe defensive burying in lateral septum of rats, and modulation by chronic treatment with desipramine. Prog Neuropsychopharmacol Biol Psychiatry 31: 482–95

Bourdelat-Parks BN, Anderson GM, Donaldson ZR, Weiss JM, Bonsall RW, et al. 2005. Effects of dopamine beta-hydroxylase genotype and disulfiram inhibition on catecholamine homeostasis in mice. Psychopharmacology (Berl) 183: 72–80

Cabib S, Castellano C, Cestari V, Filibeck U, Puglisi-Allegra S. 1991. D1 and D2 receptor antagonists differently affect cocaine-induced locomotor hyperactivity in the mouse. Psychopharmacology (Berl) 105: 335–9

Carter ME, Yizhar O, Chikahisa S, Nguyen H, Adamantidis A, et al. 2010. Tuning arousal with optogenetic modulation of locus coeruleus neurons. Nat Neurosci 13: 1526–33

Chalermpalanupap T, Kinkead B, Hu WT, Kummer MP, Hammerschmidt T, et al. 2013. Targeting norepinephrine in mild cognitive impairment and Alzheimer’s disease. Alzheimers Res Ther 5: 21

Chinen CC, Frussa-Filho R. 1999. Conditioning to injection procedures and repeated testing increase SCH 23390-induced catalepsy in mice. Neuropsychopharmacology 21: 670–8

Cho JR, Treweek JB, Robinson JE, Xiao C, Bremner LR, et al. 2017. Dorsal Raphe Dopamine Neurons Modulate Arousal and Promote Wakefulness by Salient Stimuli. Neuron 94: 1205–19 e8

Chowdhury A, Luchetti A, Fernandes G, Filho DA, Kastellakis G, et al. 2022. A locus coeruleus-dorsal CA1 dopaminergic circuit modulates memory linking. Neuron 110: 3374–88 e8

Cilz NI, Kurada L, Hu B, Lei S. 2014. Dopaminergic modulation of GABAergic transmission in the entorhinal cortex: concerted roles of alpha1 adrenoreceptors, inward rectifier K(+), and T-type Ca(2)(+) channels. Cereb Cortex 24: 3195–208

Contopoulos-Ioannidis DG, Gianniki M, Ai-Nhi Truong A, Montoya JG. 2022. Toxoplasmosis and Schizophrenia: A Systematic Review and Meta-Analysis of Prevalence and Associations and Future Directions. Psychiatr Res Clin Pract 4: 48–60

Cook TB, Brenner LA, Cloninger CR, Langenberg P, Igbide A, et al. 2015. “Latent” infection with Toxoplasma gondii: association with trait aggression and impulsivity in healthy adults. J Psychiatr Res 60: 87–94

Curtis AL, Leiser SC, Snyder K, Valentino RJ. 2012. Predator stress engages corticotropin-releasing factor and opioid systems to alter the operating mode of locus coeruleus norepinephrine neurons. Neuropharmacology 62: 1737–45

Day HE, Masini CV, Campeau S. 2004. The pattern of brain c-fos mRNA induced by a component of fox odor, 2,5-dihydro-2,4,5-trimethylthiazoline (TMT), in rats, suggests both systemic and processive stress characteristics. Brain Res 1025: 139–51

De Boer SF, Koolhaas JM. 2003. Defensive burying in rodents: ethology, neurobiology and psychopharmacology. Eur J Pharmacol 463: 145–61

De Monte FH, Canteras NS, Fernandes D, Assreuy J, Carobrez AP. 2008. New perspectives on beta-adrenergic mediation of innate and learned fear responses to predator odor. J Neurosci 28:13296–302.

Devoto P, Flore G, Saba P, Frau R, Gessa GL. 2015. Selective inhibition of dopamine-beta-hydroxylase enhances dopamine release from noradrenergic terminals in the medial prefrontal cortex. Brain Behav 5: e00393

Dias BG, Banerjee SB, Goodman JV, Ressler KJ. 2013. Towards new approaches to disorders of fear and anxiety. Curr Opin Neurobiol 23: 346–52

Dielenberg RA, McGregor IS. 2001. Defensive behavior in rats towards predatory odors: a review. Neurosci Biobehav Rev 25: 597–609

Dixit PV, Sahu R, Mishra DK. 2020. Marble-burying behavior test as a murine model of compulsive-like behavior. J Pharmacol Toxicol Methods 102: 106676

Drago F, Contarino A, Busa L. 1999. The expression of neuropeptide-induced excessive grooming behavior in dopamine D1 and D2 receptor-deficient mice. Eur J Pharmacol 365: 125–31

Dreher JK, Jackson DM. 1989. Role of D1 and D2 dopamine receptors in mediating locomotor activity elicited from the nucleus accumbens of rats. Brain Res 487: 267–77

Eban-Rothschild A, Rothschild G, Giardino WJ, Jones JR, de Lecea L. 2016. VTA dopaminergic neurons regulate ethologically relevant sleep-wake behaviors. Nat Neurosci 19: 1356–66

Feinstein DL, Kalinin S, Braun D. 2016. Causes, consequences, and cures for neuroinflammation mediated via the locus coeruleus: noradrenergic signaling system. J Neurochem 139 Suppl 2: 154–78

Fendt M, Siegl S, Steiniger-Brach B. 2005. Noradrenaline transmission within the ventral bed nucleus of the stria terminalis is critical for fear behavior induced by trimethylthiazoline, a component of fox odor. J Neurosci 25: 5998–6004

Ferrero DM, Lemon JK, Fluegge D, Pashkovski SL, Korzan WJ, et al. 2011. Detection and avoidance of a carnivore odor by prey. Proc Natl Acad Sci U S A 108: 11235–40

Fucich EA, Morilak DA. 2018. Shock-probe Defensive Burying Test to Measure Active versus Passive Coping Style in Response to an Aversive Stimulus in Rats. Bio Protoc 8

Garbe CM KE, Rawleigh JM. 1993. Novel odors evoke risk assessment and suppress appetitive behaviors in mice. Aggress Behav 19: 447–54

Garland EM, Raj SR, Peltier AC, Robertson D, Biaggioni I. 2011. A cross-sectional study contrasting olfactory function in autonomic disorders. Neurology 76: 456–60

Guild AL, Dunn AJ. 1982. Dopamine involvement in ACTH-induced grooming behavior. Pharmacol Biochem Behav 17: 31–6

Hari Dass SA, Vyas A. 2014. Toxoplasma gondii infection reduces predator aversion in rats through epigenetic modulation in the host medial amygdala. Mol Ecol 23: 6114–22

Harris NA, Isaac AT, Gunther A, Merkel K, Melchior J, et al. 2018. Dorsal BNST alpha(2A)-Adrenergic Receptors Produce HCN-Dependent Excitatory Actions That Initiate Anxiogenic Behaviors. J Neurosci 38: 8922–42

Hayley S, Borowski T, Merali Z, Anisman H. 2001. Central monoamine activity in genetically distinct strains of mice following a psychogenic stressor: effects of predator exposure. Brain Res 892: 293–300

Hu RK, Zuo Y, Ly T, Wang J, Meera P, et al. 2021. An amygdala-to-hypothalamus circuit for social reward. Nat Neurosci 24: 831–42

Hunsley MS, Palmiter RD. 2003. Norepinephrine-deficient mice exhibit normal sleep-wake states but have shorter sleep latency after mild stress and low doses of amphetamine. Sleep 26: 521–6

Hunsley MS, Palmiter RD. 2004. Altered sleep latency and arousal regulation in mice lacking norepinephrine. Pharmacol Biochem Behav 78: 765–73

Hwa LS, Neira S, Pina MM, Pati D, Calloway R, Kash TL. 2019. Predator odor increases avoidance and glutamatergic synaptic transmission in the prelimbic cortex via corticotropin-releasing factor receptor 1 signaling. Neuropsychopharmacology 44: 766–75

Iordanova MD, Yau JO, McDannald MA, Corbit LH. 2021. Neural substrates of appetitive and aversive prediction error. Neurosci Biobehav Rev 123: 337–51

Janitzky K, D’Hanis W, Krober A, Schwegler H. 2015. TMT predator odor activated neural circuit in C57BL/6J mice indicates TMT-stress as a suitable model for uncontrollable intense stress. Brain Res 1599: 1–8

Jhang J, Lee H, Kang MS, Lee HS, Park H, Han JH. 2018. Anterior cingulate cortex and its input to the basolateral amygdala control innate fear response. Nat Commun 9: 2744

Kalueff AV, Tuohimaa P. 2005. Mouse grooming microstructure is a reliable anxiety marker bidirectionally sensitive to GABAergic drugs. Eur J Pharmacol 508: 147–53

Kannan G, Moldovan K, Xiao JC, Yolken RH, Jones-Brando L, Pletnikov MV. 2010. Toxoplasma gondii strain-dependent effects on mouse behaviour. Folia Parasitol (Praha) 57: 151–5

Kedia S, Chattarji S. 2014. Marble burying as a test of the delayed anxiogenic effects of acute immobilisation stress in mice. J Neurosci Methods 233: 150–4

Kemble ED, Bolwahnn BL. 1997. Immediate and long-term effects of novel odors on risk assessment in mice. Physiol Behav 61: 543–9

Kemble ED GC, Gordon C. 1995. Effects of novel odors on intermale attack behavior in mice. Agress Behav 21: 293–99

Kempadoo KA, Mosharov EV, Choi SJ, Sulzer D, Kandel ER. 2016. Dopamine release from the locus coeruleus to the dorsal hippocampus promotes spatial learning and memory. Proc Natl Acad Sci U S A 113: 14835–40

Ketenci S, Acet NG, Saridogan GE, Aydin B, Cabadak H, Goren MZ. 2020. The Neurochemical Effects of Prazosin Treatment on Fear Circuitry in a Rat Traumatic Stress Model. Clin Psychopharmacol Neurosci 18: 219–30

Krach S, Paulus FM, Bodden M, Kircher T. 2010. The rewarding nature of social interactions. Front Behav Neurosci 4: 22

Lewis S. 2011. Autism: grooming mice to model autism. Nat Rev Neurosci 12: 248–9

Londei T, Valentini AM, Leone VG. 1998. Investigative burying by laboratory mice may involve non-functional, compulsive, behaviour. Behav Brain Res 94: 249–54

Lu J, Jhou TC, Saper CB. 2006. Identification of wake-active dopaminergic neurons in the ventral periaqueductal gray matter. J Neurosci 26: 193–202

Lustberg D, Iannitelli AF, Tillage RP, Pruitt M, Liles LC, Weinshenker D. 2020a. Central norepinephrine transmission is required for stress-induced repetitive behavior in two rodent models of obsessive-compulsive disorder. Psychopharmacology (Berl) 237: 1973–87

Lustberg D, Tillage RP, Bai Y, Pruitt M, Liles LC, Weinshenker D. 2020b. Noradrenergic circuits in the forebrain control affective responses to novelty. Psychopharmacology (Berl) 237: 3337–55

Lustberg DJ, Liu JQ, Iannitelli AF, Vanderhoof SO, Liles LC, et al. 2022. Norepinephrine and dopamine contribute to distinct repetitive behaviors induced by novel odorant stress in male and female mice. Horm Behav 144: 105205

McCall JG, Al-Hasani R, Siuda ER, Hong DY, Norris AJ, et al. 2015. CRH Engagement of the Locus Coeruleus Noradrenergic System Mediates Stress-Induced Anxiety. Neuron 87: 605–20

McGregor IS, Hargreaves GA, Apfelbach R, Hunt GE. 2004. Neural correlates of cat odor-induced anxiety in rats: region-specific effects of the benzodiazepine midazolam. J Neurosci 24: 4134–44

McGregor IS, Schrama L, Ambermoon P, Dielenberg RA. 2002. Not all ‘predator odours’ are equal: cat odour but not 2,4,5 trimethylthiazoline (TMT; fox odour) elicits specific defensive behaviours in rats. Behav Brain Res 129: 1–16

Miller SM, Marcotulli D, Shen A, Zweifel LS. 2019. Divergent medial amygdala projections regulate approach-avoidance conflict behavior. Nat Neurosci 22: 565–75

Mitchell HA, Bogenpohl JW, Liles LC, Epstein MP, Bozyczko-Coyne D, et al. 2008. Behavioral responses of dopamine beta-hydroxylase knockout mice to modafinil suggest a dual noradrenergic-dopaminergic mechanism of action. Pharmacol Biochem Behav 91: 217–22

Mitchell HA, Weinshenker D. 2010. Good night and good luck: norepinephrine in sleep pharmacology. Biochem Pharmacol 79: 801–9

Nayeri T, Sarvi S, Daryani A. 2022. Toxoplasmosis: Targeting neurotransmitter systems in psychiatric disorders. Metab Brain Dis 37: 123–46

Nayeri T, Sarvi S, Moosazadeh M, Hosseininejad Z, Sharif M, et al. 2020. Relationship between toxoplasmosis and autism: A systematic review and meta-analysis. Microb Pathog 147: 104434

Nordstrom EJ, Burton FH. 2002. A transgenic model of comorbid Tourette’s syndrome and obsessive-compulsive disorder circuitry. Mol Psychiatry 7: 617–25, 524

Ohkubo N, Aoto M, Kon K, Mitsuda N. 2019. Lack of zinc finger protein 521 upregulates dopamine beta-hydroxylase expression in the mouse brain, leading to abnormal behavior. Life Sci 231: 116559

Ozkan M, Johnson NW, Sehirli US, Woodhall GL, Stanford IM. 2017. Dopamine acting at D1-like, D2-like and alpha1-adrenergic receptors differentially modulates theta and gamma oscillatory activity in primary motor cortex. PLoS One 12: e0181633

Paladini CA, Beckstead MJ, Weinshenker D. 2007. Electrophysiological properties of catecholaminergic neurons in the norepinephrine-deficient mouse. Neuroscience 144: 1067–74

Paladini CA, Fiorillo CD, Morikawa H, Williams JT. 2001. Amphetamine selectively blocks inhibitory glutamate transmission in dopamine neurons. Nat Neurosci 4: 275–81

Petter EA, Fallon IP, Hughes RN, Watson GDR, Meck WH, et al. 2023. Elucidating a locus coeruleus-dentate gyrus dopamine pathway for operant reinforcement. Elife 12

Pina MM, Cunningham CL. 2014. Effects of dopamine receptor antagonists on the acquisition of ethanol-induced conditioned place preference in mice. Psychopharmacology (Berl) 231: 459–68

Poe GR, Foote S, Eschenko O, Johansen JP, Bouret S, et al. 2020. Locus coeruleus: a new look at the blue spot. Nat Rev Neurosci 21: 644–59

Pond HL, Heller AT, Gural BM, McKissick OP, Wilkinson MK, Manzini MC. 2021. Digging behavior discrimination test to probe burrowing and exploratory digging in male and female mice. J Neurosci Res 99: 2046–58

Porter-Stransky KA, Centanni SW, Karne SL, Odil LM, Fekir S, et al. 2019. Noradrenergic Transmission at Alpha1-Adrenergic Receptors in the Ventral Periaqueductal Gray Modulates Arousal. Biol Psychiatry 85: 237–47

Rosen JB, Pagani JH, Rolla KL, Davis C. 2008. Analysis of behavioral constraints and the neuroanatomy of fear to the predator odor trimethylthiazoline: a model for animal phobias. Neurosci Biobehav Rev 32: 1267–76

Sara SJ. 2009. The locus coeruleus and noradrenergic modulation of cognition. Nat Rev Neurosci 10: 211–23

Schank JR, Liles LC, Weinshenker D. 2008. Norepinephrine signaling through beta-adrenergic receptors is critical for expression of cocaine-induced anxiety. Biol Psychiatry 63: 1007–12

Schroeder JP, Cooper DA, Schank JR, Lyle MA, Gaval-Cruz M, et al. 2010. Disulfiram attenuates drug-primed reinstatement of cocaine seeking via inhibition of dopamine beta-hydroxylase. Neuropsychopharmacology 35: 2440–9

Schwarz LA, Luo L. 2015. Organization of the locus coeruleus-norepinephrine system. Curr Biol 25: R1051–R56

Sluyter F, Korte SM, Bohus B, Van Oortmerssen GA. 1996. Behavioral stress response of genetically selected aggressive and nonaggressive wild house mice in the shock-probe/defensive burying test. Pharmacol Biochem Behav 54: 113–6

Smolinsky AN BC, LaPorte JL, Kalueff AV. 2009. Analysis of grooming behavior and its utility in studying animal stress, anxiety, and depression. pp. 21–36. Springer.

Stanley WC, Li B, Bonhaus DW, Johnson LG, Lee K, et al. 1997. Catecholamine modulatory effects of nepicastat (RS-25560-197), a novel, potent and selective inhibitor of dopamine-beta-hydroxylase. Br J Pharmacol 121: 1803–9

Stone EA, Yan L, Ahsan MR, Lehmann ML, Yeretsian J, Quartermain D. 2006. Role of CNS alpha1-adrenoceptor activity in central fos responses to novelty. Synapse 59: 299–307

Stone EA, Zhang Y, Rosengarten H, Yeretsian J, Quartermain D. 1999. Brain alpha 1-adrenergic neurotransmission is necessary for behavioral activation to environmental change in mice. Neuroscience 94: 1245–52

Szot P, Weinshenker D, White SS, Robbins CA, Rust NC, et al. 1999. Norepinephrine-deficient mice have increased susceptibility to seizure-inducing stimuli. J Neurosci 19: 10985–92

Takahashi LK. 2014. Olfactory systems and neural circuits that modulate predator odor fear. Front Behav Neurosci 8: 72

Takeuchi T, Duszkiewicz AJ, Sonneborn A, Spooner PA, Yamasaki M, et al. 2016. Locus coeruleus and dopaminergic consolidation of everyday memory. Nature 537: 357–62

Tedford E, Badya NB, Laing C, Asaoka N, Kaneko S, et al. 2023. Infection-induced extracellular vesicles evoke neuronal transcriptional and epigenetic changes. Sci Rep 13: 6913

Thomas SA, Marck BT, Palmiter RD, Matsumoto AM. 1998. Restoration of norepinephrine and reversal of phenotypes in mice lacking dopamine beta-hydroxylase. J Neurochem 70: 2468–76

Thomas SA, Matsumoto AM, Palmiter RD. 1995. Noradrenaline is essential for mouse fetal development. Nature 374: 643–6

Thomas SA, Palmiter RD. 1997. Impaired maternal behavior in mice lacking norepinephrine and epinephrine. Cell 91: 583–92

Tillage RP, Sciolino NR, Plummer NW, Lustberg D, Liles LC, et al. 2020. Elimination of galanin synthesis in noradrenergic neurons reduces galanin in select brain areas and promotes active coping behaviors. Brain Struct Funct 225: 785–803

Timmermans PB, van Zwieten PA. 1982. alpha 2 adrenoceptors: classification, localization, mechanisms, and targets for drugs. J Med Chem 25: 1389–401

Torrey EF, Yolken RH. 2003. Toxoplasma gondii and schizophrenia. Emerg Infect Dis 9: 1375–80

Treit D, Pinel JP, Fibiger HC. 1981. Conditioned defensive burying: a new paradigm for the study of anxiolytic agents. Pharmacol Biochem Behav 15: 619–26

Vincenz D, Wernecke KEA, Fendt M, Goldschmidt J. 2017. Habenula and interpeduncular nucleus differentially modulate predator odor-induced innate fear behavior in rats. Behav Brain Res 332: 164–71

Virus MA, Ehrhorn EG, Lui LM, Davis PH. 2021. Neurological and Neurobehavioral Disorders Associated with Toxoplasma gondii Infection in Humans. J Parasitol Res 2021: 6634807

Vyas A, Kim SK, Giacomini N, Boothroyd JC, Sapolsky RM. 2007. Behavioral changes induced by Toxoplasma infection of rodents are highly specific to aversion of cat odors. Proc Natl Acad Sci U S A 104: 6442–7

Wagatsuma A, Okuyama T, Sun C, Smith LM, Abe K, Tonegawa S. 2018. Locus coeruleus input to hippocampal CA3 drives single-trial learning of a novel context. Proc Natl Acad Sci U S A 115: E310–E16

Wang M, Ramos BP, Paspalas CD, Shu Y, Simen A, et al. 2007. Alpha2A-adrenoceptors strengthen working memory networks by inhibiting cAMP-HCN channel signaling in prefrontal cortex. Cell 129: 397–410

Webster JP, Brunton CF, MacDonald DW. 1994. Effect of Toxoplasma gondii upon neophobic behaviour in wild brown rats, Rattus norvegicus. Parasitology 109 ( Pt 1): 37–43

Weinshenker D, Miller NS, Blizinsky K, Laughlin ML, Palmiter RD. 2002. Mice with chronic norepinephrine deficiency resemble amphetamine-sensitized animals. Proc Natl Acad Sci U S A 99: 13873–7

Welch JM, Lu J, Rodriguiz RM, Trotta NC, Peca J, et al. 2007. Cortico-striatal synaptic defects and OCD-like behaviours in Sapap3-mutant mice. Nature 448: 894–900

Xu HY, Liu YJ, Xu MY, Zhang YH, Zhang JX, Wu YJ. 2012. Inactivation of the bed nucleus of the stria terminalis suppresses the innate fear responses of rats induced by the odor of cat urine. Neuroscience 221: 21–7

Yamaguchi H, Hopf FW, Li SB, de Lecea L. 2018. In vivo cell type-specific CRISPR knockdown of dopamine beta hydroxylase reduces locus coeruleus evoked wakefulness. Nat Commun 9: 5211

Yan T, Xu M, Wu B, Liao Z, Liu Z, et al. 2016. The effect of Schisandra chinensis extracts on depression by noradrenergic, dopaminergic, GABAergic and glutamatergic systems in the forced swim test in mice. Food Funct 7: 2811–9

